# Nonclinical pharmacokinetics and relative efficacy of the first 25 novel tuberculosis drug combinations from the PAN-TB consortium: Use of the BALB/c relapsing mouse model and combination pharmacokinetics within a modeling-based framework

**DOI:** 10.64898/2026.05.05.722941

**Authors:** Sylvie Sordello, Alessia Tagliavini, Xavier Boulenc, Laure Brock, Simone Zannoni, Chiara Roversi, Roberto Visentin, Darren Metcalf, Denise Federico, Simone Modolo, Giulia Calusi, Roberto Petterlini, Guillaume Golovkine, Cécile Pascal, Emilie Huc Claustre, Zoï Vahlas, Marco Pergher, Khisimuzi Mdluli, Micha Levi, Todd A. Black, Robert H. Bates, David R Willé, Yongge Liu, Yohei Hayashi, Clara Aguilar-Pérez, David Hermann, Debra Flood, Anna M. Upton

## Abstract

The Project to Accelerate New Treatments for Tuberculosis (PAN-TB) aims to accelerate development of shorter, simpler and safer pan-TB combinations, effective for use in both Drug Susceptible (DS)- and Drug Resistant (DR)- TB patients. Towards this aim, bactericidal and sterilizing activity of 25 priority 4-drug combinations was evaluated at doses targeting clinically relevant exposures, in the BALB/c relapsing mouse model of TB. The combinations comprised 8 PAN-TB drugs and candidates: bedaquiline (B), pretomanid (Pa), delamanid (Del), quabodepistat (Q), sutezolid (Sut), GSK2556286 (286), GSK3211830 (830) and ganfeborole (GSK3036656, (656)). Combination PK studies in infected mice enabled dose selection and a population-PK approach guided dosing so that compounds should achieve mean AUC_0-24_ within 2-fold of their clinical target exposures during the efficacy studies. All test combinations showed time-dependent bactericidal activity, with six regimens reducing lung bacterial burdens below the limit of detection with 8 weeks’ treatment, similar to the comparator BPaMZ (M is moxifloxacin and Z as pyrazinamide). Cure/Relapse data were modelled to derive population time to cure 90% mice (T90) values. Fifteen PAN-TB combinations had T90s of less than 5 months, sterilizing mice faster than the standard of care for drug susceptible TB, RHZE/RH. The best-performing PAN-TB combinations, BPa830Sut, BPa286Sut and BQSut286, cured 90% of mice in less than 3 months. These 3 top-ranked 4-drug combinations are all centered on a diarylquinoline (B)/oxazolidinone (Sut) core, together with the nitroimidazole (Pa) or a DprE1 inhibitor (Q) plus a novel agent such as the LeuRS inhibitor (830) or the Rv1625c agonist (286).

## INTRODUCTION

Tuberculosis (TB) remains a global health crisis with 1.25 million deaths reported in 2023 [1]. Available treatments are long, complex and poorly tolerated, resulting in suboptimal adherence which impacts patient outcomes. Due to the presence of significant drug-resistance, access to rapid drug susceptibility testing (DST) is critical to appropriate treatment; the currently limited access presents another barrier to TB control [2]. Development of novel pan-TB combinations, defined by World Health Organization (WHO) as first-line regimens that could be used without prior knowledge of a TB patient’s drug-resistance profile [3], would obviate the need for DST, eliminating the time gap between diagnosis and start of effective treatment. To maximize benefits, pan-TB regimens should be shorter, simpler, and safer than existing treatments. Introducing such regimens in high TB burden countries could result in substantial health improvements as well as savings to patients and health systems [4]. Although progress has been made towards treatments that fit WHO target regimen profiles for drug-susceptible TB (DS-TB) and rifampicin resistant TB (RR-TB) [3], more limited progress has been made towards development of pan-TB regimens.

The Project to Accelerate New Treatments for Tuberculosis (PAN-TB) is a philanthropic, non-profit and private sector collaboration. The Collaboration aims to accelerate prioritization and development of promising pan-TB drug combinations by leveraging assets, resources, technologies and expertise of its members ([Tuberculosis Prevention](https://www.pan-tb.org/) | PAN-TB). At the outset of the Collaboration, eight candidates and marketed drugs were identified among the members as PAN-TB priorities, due to their pan-TB potential (i.e. no or limited existing clinical resistance). These comprised bedaquiline (B), an ATP synthase inhibitor - [5], delamanid (Del) and pretomanid (Pa), two nitroimidazole mycolic acid synthesis inhibitors [6–7], quabodepistat (Q) an inhibitor of decaprenylphosphoryl-β-D-ribose 2’-epimerase [DprE1] [8], GSK2556286 (286) an Rv1625c agonist and, c-AMP mediated cholesterol catabolism inhibitor [10], sutezolid (Sut) an RNA translation inhibitor [11], GSK3211830 (830) and ganfeborole (formerly GSK3036656, listed here as 656), leucyl-tRNA synthetase (LeuRS) inhibitors [9]. These two LeuRS inhibitors are close analogues with very similar PK and efficacy data. The consortium initially focused on testing 830 but later shifted toward 656 as it advanced further in the clinic.

The Collaboration focuses on 4-drug pan-TB combinations, hypothesizing that treatment with 4 drugs with novel and differing modes of action, will maximize efficacy and minimize emergence of resistance. Towards prioritizing the most promising 4-drug combinations comprised of the initial 8 PAN-TB candidates, the Collaboration selected 25 unique regimens, avoiding inclusion of two nitroimidazoles in the same combination, for nonclinical efficacy assessment. Some of the candidates had previously demonstrated bactericidal efficacy and treatment-shortening activity when evaluated within TB drug combinations in the well-established BALB/c relapsing mouse model of TB (RMM) [8, 12, 13, 14, 15, 16]. None had been evaluated within the context of these specific 25 combinations.

This work evaluated and compared the bactericidal and sterilizing efficacy of the 25 drug combinations in the BALB/c RMM of TB, at exposures relevant to clinical efficacy targets. An additional objective was to compare their estimated RMM time to cure parameters to those of clinical benchmark regimens. Benchmark TB combinations used in the reported RMM studies included the standard of care for drug-susceptible TB, Rifampicin (R), Isoniazid (H), Pyrazinamide (Z), Ethambutol given as RHZE/RH and BPaMZ. BPaMZ is a regimen that has been evaluated in clinical trials and demonstrated time to culture conversion that was more rapid than RHZE/RH [17]. However, the regimen did not meet the key secondary efficacy endpoint due to adverse events resulting in treatment withdrawal and additionally, includes pyrazinamide and moxifloxacin, to which there is significant background resistance. Therefore, BPaMZ cannot be considered as a potential pan-TB regimen. We sought to identify pan-TB regimens with efficacy superior to BPaMZ in this well-established mouse TB model.

## RESULTS

### Combination Pharmacokinetics Performed in Infected Mice Enabled Accurate Selection of Human Equivalent Doses

The 25 PAN-TB test combinations are listed in **Table 1**. Doses for evaluation in RMM efficacy studies were selected with the aim of achieving steady-state area under the concentration time curve from 0-24 hours (AUC_0-24_) for each drug, across all test combinations, within 2-fold of observed or targeted clinical plasma concentrations (**Table 2).** Preliminary dose selection was based on available mouse single drug plasma PK data and rodent tolerability data. Doses were refined based on data generated through 5-day repeat-dose combination PK studies, conducted for each test combination, administered at the preliminary doses to *M.tb*-infected mice. The average plasma AUC_0-24_ observed for each drug, across the relevant 4-drug combinations in mice, was compared to the corresponding observed or target clinical value. As a result of these comparisons, the final dose for use in efficacy evaluations was decreased for Del, and increased for both 830 and 656, compared to their preliminary doses (**Table 2**). Q exposure was variable between tested combinations. Although the median Q exposure in mice, at the preliminary dose, was higher than the clinical target, the final dose selected was slightly higher than the preliminary, to ensure exposures across all test combinations were no more than 2-fold lower than the clinical target.

**Table 1.**
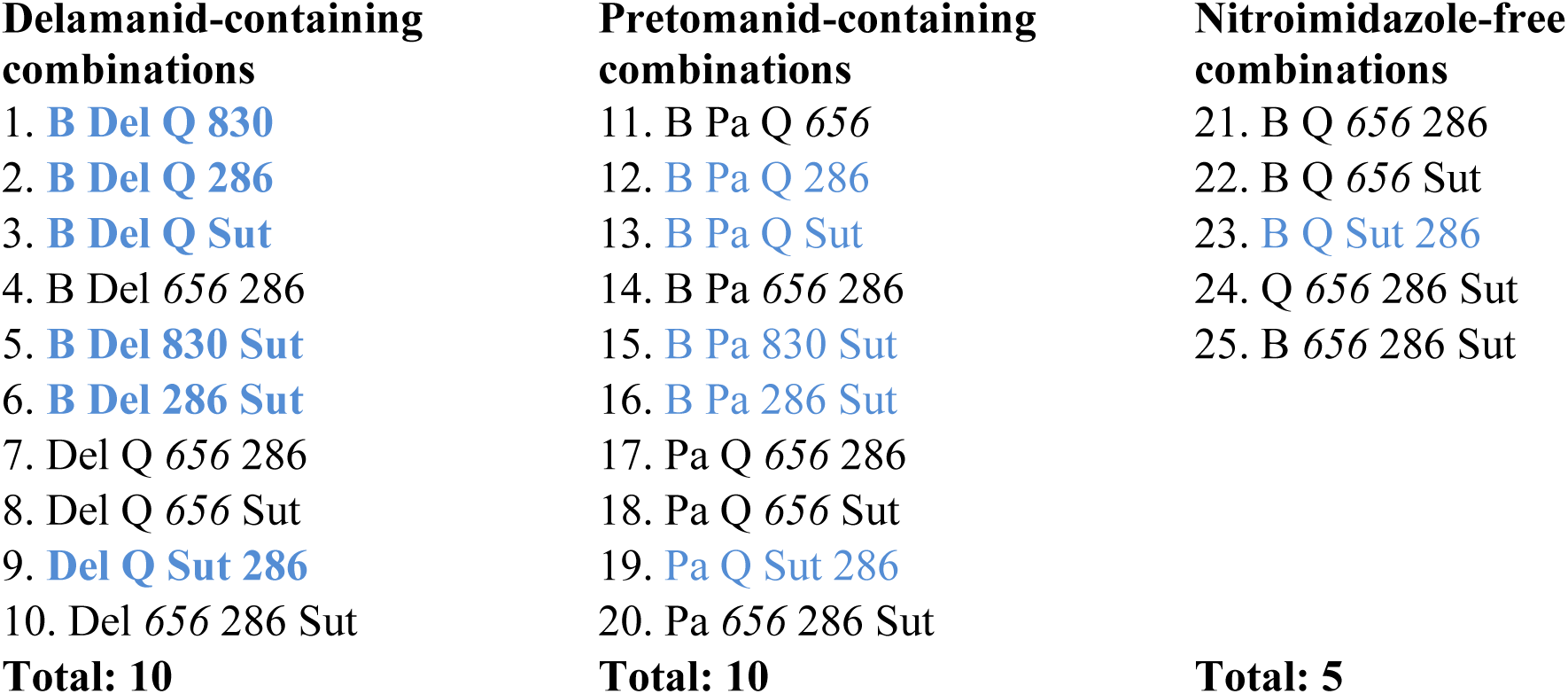
PAN-TB 4-drug Combinations. Combinations in blue font were tested in Study 1. Combinations in black font were tested in Study 2. 830 was replaced by 656 in Study 2.

**Table 2.**
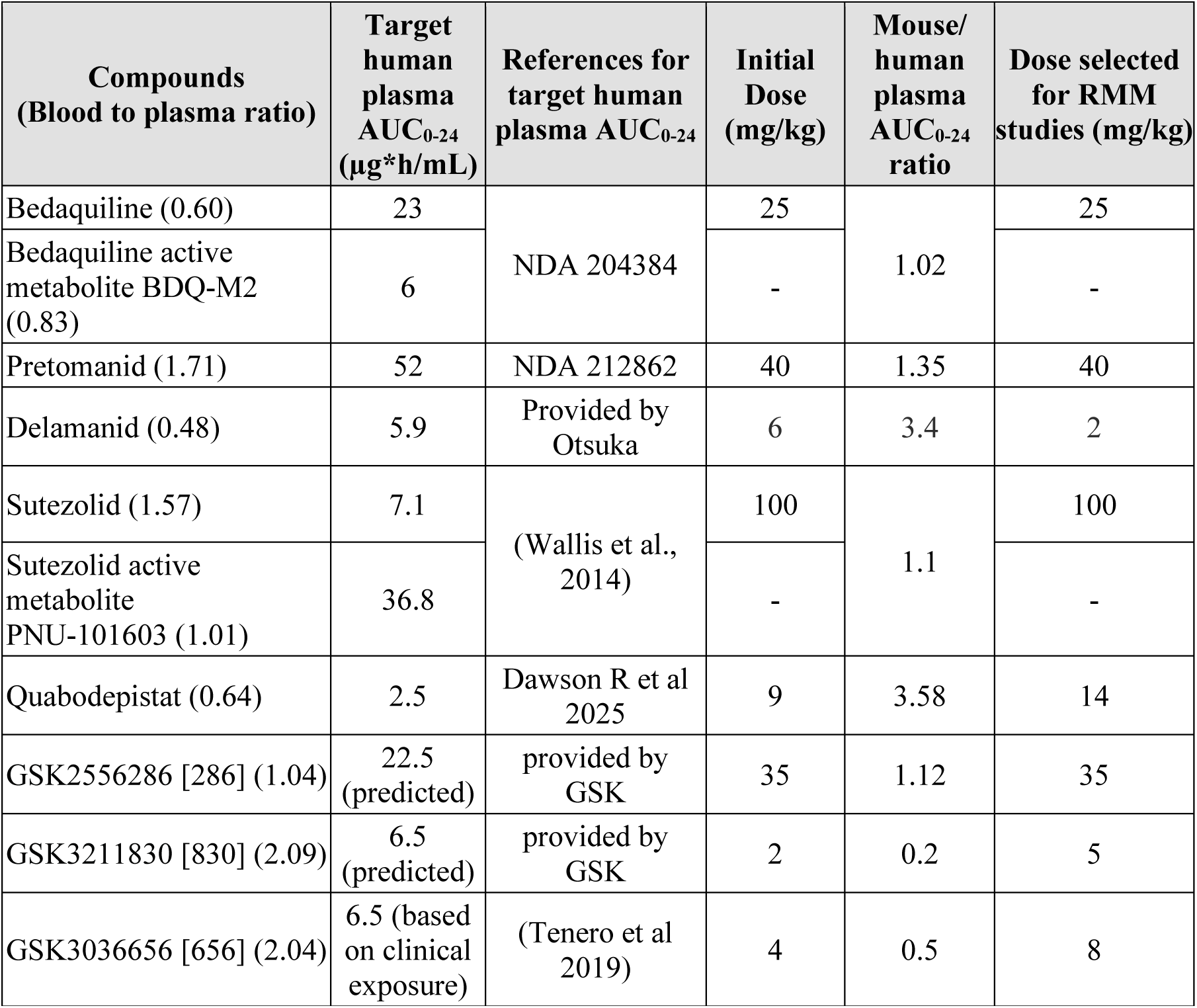
Doses Used in Mouse Studies to reach Clinical Target Plasma Exposures.

| Compounds<br>(Blood to plasma ratio) | Target<br>human<br>plasma<br>AUC<sub>0-24</sub><br>(µg*h/mL) | References for<br>target human<br>plasma AUC<sub>0-24</sub> | Initial<br>Dose<br>(mg/kg) | Mouse/<br>human<br>plasma<br>AUC<sub>0-24</sub><br>ratio | Dose selected<br>for RMM<br>studies (mg/kg) |
| --- | --- | --- | --- | --- | --- |
| Bedaquiline (0.60) | 23 | NDA 204384 | 25 | 1.02 | 25 |
| Bedaquiline active<br>metabolite BDQ-M2<br>(0.83) | 6 |  | - |  | - |
| Pretomanid (1.71) | 52 | NDA 212862 | 40 | 1.35 | 40 |
| Delamanid (0.48) | 5.9 | Provided by<br>Otsuka | 6 | 3.4 | 2 |
| Sutezolid (1.57) | 7.1 | (Wallis et al.,<br>2014) | 100 | 1.1 | 100 |
| Sutezolid active<br>metabolite<br>PNU-101603 (1.01) | 36.8 |  | - |  | - |
| Quabodepistat (0.64) | 2.5 | Dawson R et al<br>2025 | 9 | 3.58 | 14 |
| GSK2556286 [286] (1.04) | 22.5<br>(predicted) | provided by<br>GSK | 35 | 1.12 | 35 |
| GSK3211830 [830] (2.09) | 6.5<br>(predicted) | provided by<br>GSK | 2 | 0.2 | 5 |
| GSK3036656 [656] (2.04) | 6.5 (based<br>on clinical<br>exposure) | (Tenero et al<br>2019) | 4 | 0.5 | 8 |

### RMM plasma exposures were within 2-fold of their clinical targets for most combination components

To evaluate PK exposure achieved during RMM efficacy studies for each drug across the test combinations, population PK (popPK) analysis was performed using sparse-sampled drug concentration data obtained after 8 weeks’ dosing during the RMM studies, together with data generated through the combination PK studies. Individual RMM drug exposure distributions across all tested combinations are shown in **Figures S1-S8 and Table S2**. For most of the tested compounds, i.e., 656, B, Del, Pa and Sut, their overall distributions of RMM study drug exposures (derived plasma AUC_0-24_ values) achieved after 8 weeks’ dosing were within 2-fold of their clinical targets (**Figure 1 and Table S2**). The Q RMM exposures were variable across the tested combinations: none were more than 2-fold below the targeted exposure, and while the median observed Q plasma AUC_0-24_ was slightly more than 2-fold higher than the clinical target, exposures were within 2-fold of the clinical target for 8 out of 16 of the tested Q-containing combinations, namely when Q was co-administered with B and in the absence of Pa (**Figure S7).** The 286 RMM exposures were slightly more than 2-fold lower than its clinical target on average **(Figure 1** and **Table S2)**. Notably, AUC_0-24_ values for 286 were lowest when co-administered with B (**Figure S1**).

**Figure 1.** Distribution of plasma AUC_0-24_ (AUC_0-24/MIC_ for B and Sut) from popPK analysis versus clinical target.

Finally, 830 RMM exposures were more than 2-fold lower than the clinical target in all tested combinations (**Figure S3**). When comparing exposures of companion drugs between Del- and Pa- containing regimens, B exposure is, at least two-times higher in the Pa-containing regimens than those with Del, except for the regimens of BDel656286 and BPa656286, in which B exposures are similar (**Figure S4**). Similarly, Q exposure is more than two-times higher in BPaQX regimens than those in the BDelQX regimens (**Figure S7**)

### Six PAN-TB drug combinations demonstrated bactericidal activity in mice that was at least as rapid as BPaMZ

All 25 4-drug combinations demonstrated time-dependent bactericidal activity over the course of 8 weeks’ treatment (**Figures 2A and 2B**). BPa286Sut and BQSut286 reduced lung CFU to undetectable levels following 4 weeks of treatment, whereas BDel286Sut, BDel830Sut, BPa830Sut, BPaQSut and BPaMZ required 8 weeks’ treatment. For the other 19 test combinations, as for the benchmark RHZE/RH, colonies were still detectable after 8-weeks’ treatment (**Figures 2A and 2B**). However, 15 of these 19 reduced the mouse lung bacterial burden (CFU/ lung) to a greater extent than RHZE/RH by the 8 week treatment timepoint: BPa656286, BQ656286, BQ656Sut, BDelQSut, BDelQ286, BDelQ830, BDel656286, PaQSut286, PaQ656Sut reduced bacterial burden by an additional 2Log_10,_ and BDelQ830, BPaQ286, B656286Sut, PaQ656286, Del656286Sut, BPaQ656 reduced CFU/ lung by an additional 1Log_10_ compared to RHZE/RH (**Tables 3A and 3B**).

**Figure 2.** Lung bacterial burden at end of treatment in Study 1 (A) and Study 2 (B). Whole lung CFU of BALB/c mice, intranasally infected with *M.tb* H37Rv, after different durations of oral treatment with 4-drug combinations, dosed 5/7. Treatments were initiated 2-weeks post infection.

**Table 3:**
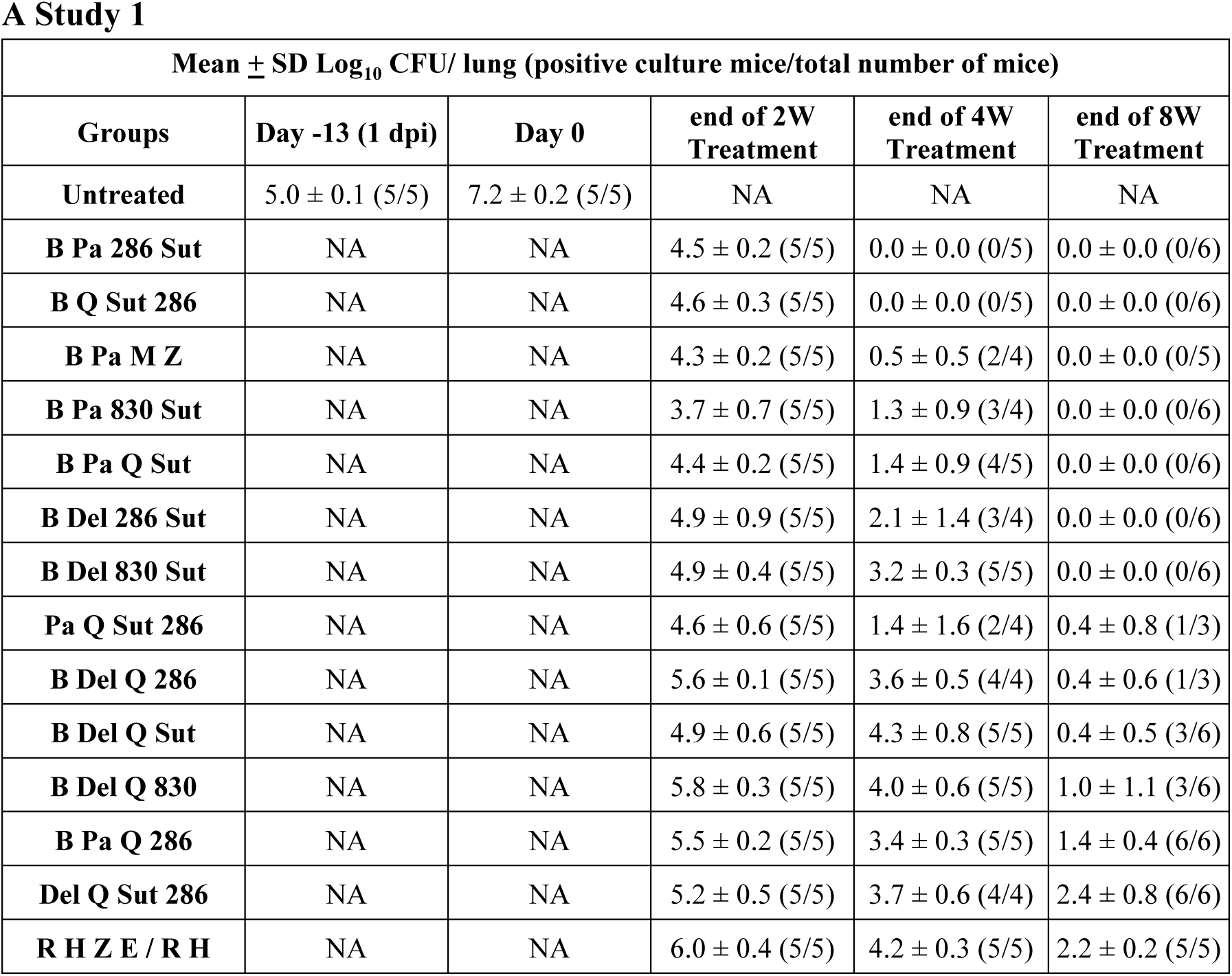
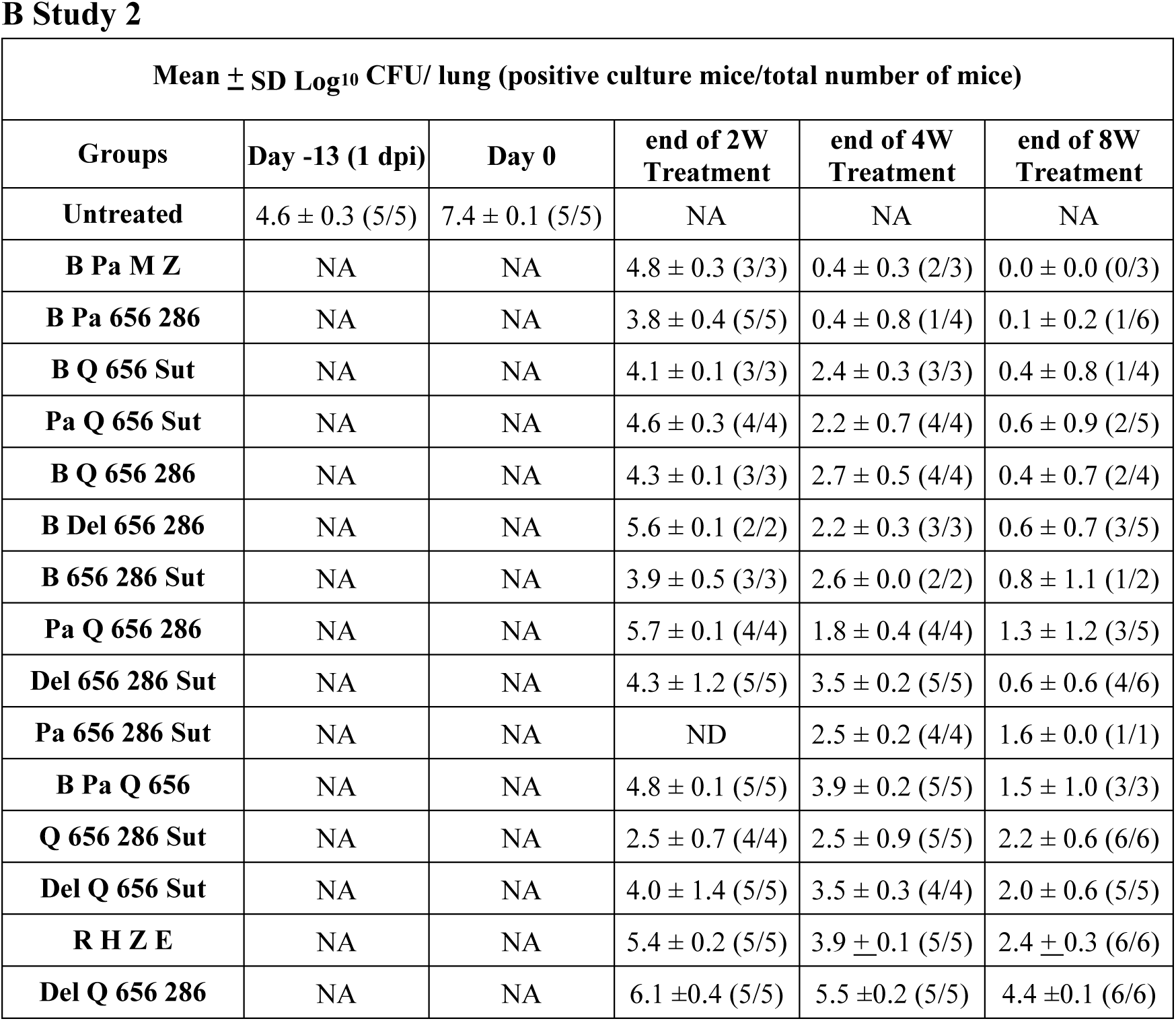
Lung bacterial burden prior and at the end of treatment in Study 1 (A) and Study 2 (B) after different durations of oral treatment with 4-drug combinations.

**A Study 1**
| **Mean ± SD Log<sub>10</sub> CFU/ lung (positive culture mice/total number of mice)** |  |  |  |  |  |
| --- | --- | --- | --- | --- | --- |
| **Groups** | **Day -13 (1 dpi)** | **Day 0** | **end of 2W Treatment** | **end of 4W Treatment** | **end of 8W Treatment** |
| **Untreated** | 5.0 ± 0.1 (5/5) | 7.2 ± 0.2 (5/5) | NA | NA | NA |
| **B Pa 286 Sut** | NA | NA | 4.5 ± 0.2 (5/5) | 0.0 ± 0.0 (0/5) | 0.0 ± 0.0 (0/6) |
| **B Q Sut 286** | NA | NA | 4.6 ± 0.3 (5/5) | 0.0 ± 0.0 (0/5) | 0.0 ± 0.0 (0/6) |
| **B Pa M Z** | NA | NA | 4.3 ± 0.2 (5/5) | 0.5 ± 0.5 (2/4) | 0.0 ± 0.0 (0/5) |
| **B Pa 830 Sut** | NA | NA | 3.7 ± 0.7 (5/5) | 1.3 ± 0.9 (3/4) | 0.0 ± 0.0 (0/6) |
| **B Pa Q Sut** | NA | NA | 4.4 ± 0.2 (5/5) | 1.4 ± 0.9 (4/5) | 0.0 ± 0.0 (0/6) |
| **B Del 286 Sut** | NA | NA | 4.9 ± 0.9 (5/5) | 2.1 ± 1.4 (3/4) | 0.0 ± 0.0 (0/6) |
| **B Del 830 Sut** | NA | NA | 4.9 ± 0.4 (5/5) | 3.2 ± 0.3 (5/5) | 0.0 ± 0.0 (0/6) |
| **Pa Q Sut 286** | NA | NA | 4.6 ± 0.6 (5/5) | 1.4 ± 1.6 (2/4) | 0.4 ± 0.8 (1/3) |
| **B Del Q 286** | NA | NA | 5.6 ± 0.1 (5/5) | 3.6 ± 0.5 (4/4) | 0.4 ± 0.6 (1/3) |
| **B Del Q Sut** | NA | NA | 4.9 ± 0.6 (5/5) | 4.3 ± 0.8 (5/5) | 0.4 ± 0.5 (3/6) |
| **B Del Q 830** | NA | NA | 5.8 ± 0.3 (5/5) | 4.0 ± 0.6 (5/5) | 1.0 ± 1.1 (3/6) |
| **B Pa Q 286** | NA | NA | 5.5 ± 0.2 (5/5) | 3.4 ± 0.3 (5/5) | 1.4 ± 0.4 (6/6) |
| **Del Q Sut 286** | NA | NA | 5.2 ± 0.5 (5/5) | 3.7 ± 0.6 (4/4) | 2.4 ± 0.8 (6/6) |
| **R H Z E / R H** | NA | NA | 6.0 ± 0.4 (5/5) | 4.2 ± 0.3 (5/5) | 2.2 ± 0.2 (5/5) |

**Table 3:** Lung bacterial burden prior and at the end of treatment in Study 1 (A) and Study 866 2 (B) after different durations of oral treatment with 4-drug combinations.
| Mean ± SD Log<sub>10</sub> CFU/ lung (positive culture mice/total number of mice) | | | | | |
| --- | --- | --- | --- | --- | --- |
| Groups | Day -13 (1 dpi) | Day 0 | end of 2W Treatment | end of 4W Treatment | end of 8W Treatment |
| Untreated | 4.6 ± 0.3 (5/5) | 7.4 ± 0.1 (5/5) | NA | NA | NA |
| B Pa M Z | NA | NA | 4.8 ± 0.3 (3/3) | 0.4 ± 0.3 (2/3) | 0.0 ± 0.0 (0/3) |
| B Pa 656 286 | NA | NA | 3.8 ± 0.4 (5/5) | 0.4 ± 0.8 (1/4) | 0.1 ± 0.2 (1/6) |
| B Q 656 Sut | NA | NA | 4.1 ± 0.1 (3/3) | 2.4 ± 0.3 (3/3) | 0.4 ± 0.8 (1/4) |
| Pa Q 656 Sut | NA | NA | 4.6 ± 0.3 (4/4) | 2.2 ± 0.7 (4/4) | 0.6 ± 0.9 (2/5) |
| B Q 656 286 | NA | NA | 4.3 ± 0.1 (3/3) | 2.7 ± 0.5 (4/4) | 0.4 ± 0.7 (2/4) |
| B Del 656 286 | NA | NA | 5.6 ± 0.1 (2/2) | 2.2 ± 0.3 (3/3) | 0.6 ± 0.7 (3/5) |
| B 656 286 Sut | NA | NA | 3.9 ± 0.5 (3/3) | 2.6 ± 0.0 (2/2) | 0.8 ± 1.1 (1/2) |
| Pa Q 656 286 | NA | NA | 5.7 ± 0.1 (4/4) | 1.8 ± 0.4 (4/4) | 1.3 ± 1.2 (3/5) |
| Del 656 286 Sut | NA | NA | 4.3 ± 1.2 (5/5) | 3.5 ± 0.2 (5/5) | 0.6 ± 0.6 (4/6) |
| Pa 656 286 Sut | NA | NA | ND | 2.5 ± 0.2 (4/4) | 1.6 ± 0.0 (1/1) |
| B Pa Q 656 | NA | NA | 4.8 ± 0.1 (5/5) | 3.9 ± 0.2 (5/5) | 1.5 ± 1.0 (3/3) |
| Q 656 286 Sut | NA | NA | 2.5 ± 0.7 (4/4) | 2.5 ± 0.9 (5/5) | 2.2 ± 0.6 (6/6) |
| Del Q 656 Sut | NA | NA | 4.0 ± 1.4 (5/5) | 3.5 ± 0.3 (4/4) | 2.0 ± 0.6 (5/5) |
| R H Z E | NA | NA | 5.4 ± 0.2 (5/5) | 3.9 ± 0.1 (5/5) | 2.4 ± 0.3 (6/6) |
| Del Q 656 286 | NA | NA | 6.1 ± 0.4 (5/5) | 5.5 ± 0.2 (5/5) | 4.4 ± 0.1 (6/6) |

### Three PAN-TB 4-drug combinations cured 90% of mice in less than 3 months, out-performing RHZE/RH but not BPaMZ

To establish the relationship between the tested regimens and length of treatment required to achieve a durable cure in mice, we quantified the proportion of mice that were culture positive twelve weeks after the end of treatment (i.e. exhibiting relapse) (**Tables 4A and 4B)**. No relapse events were recorded for any BPaMZ-treated mice after 6 weeks’ treatment. In contrast, at least 16 weeks’ treatment were required to achieve 100% cure (no relapse) for RHZE/RH. Of the PAN-TB test combinations, the best-performing were BPa830Sut (100% mice cured after 8 weeks’ treatment) and BPa286Sut, which cured all mice after 10 weeks’ treatment. Six additional test combinations demonstrated 100% cure after 12 weeks of treatment (**Table 4A**). On the other hand, 8 of the 25 test combinations failed to effect cure after 12 weeks’ treatment: all mice had detectable colonies (relapse) 12 weeks after treatment cessation (**Table 4A and B**).

**Table 4.**
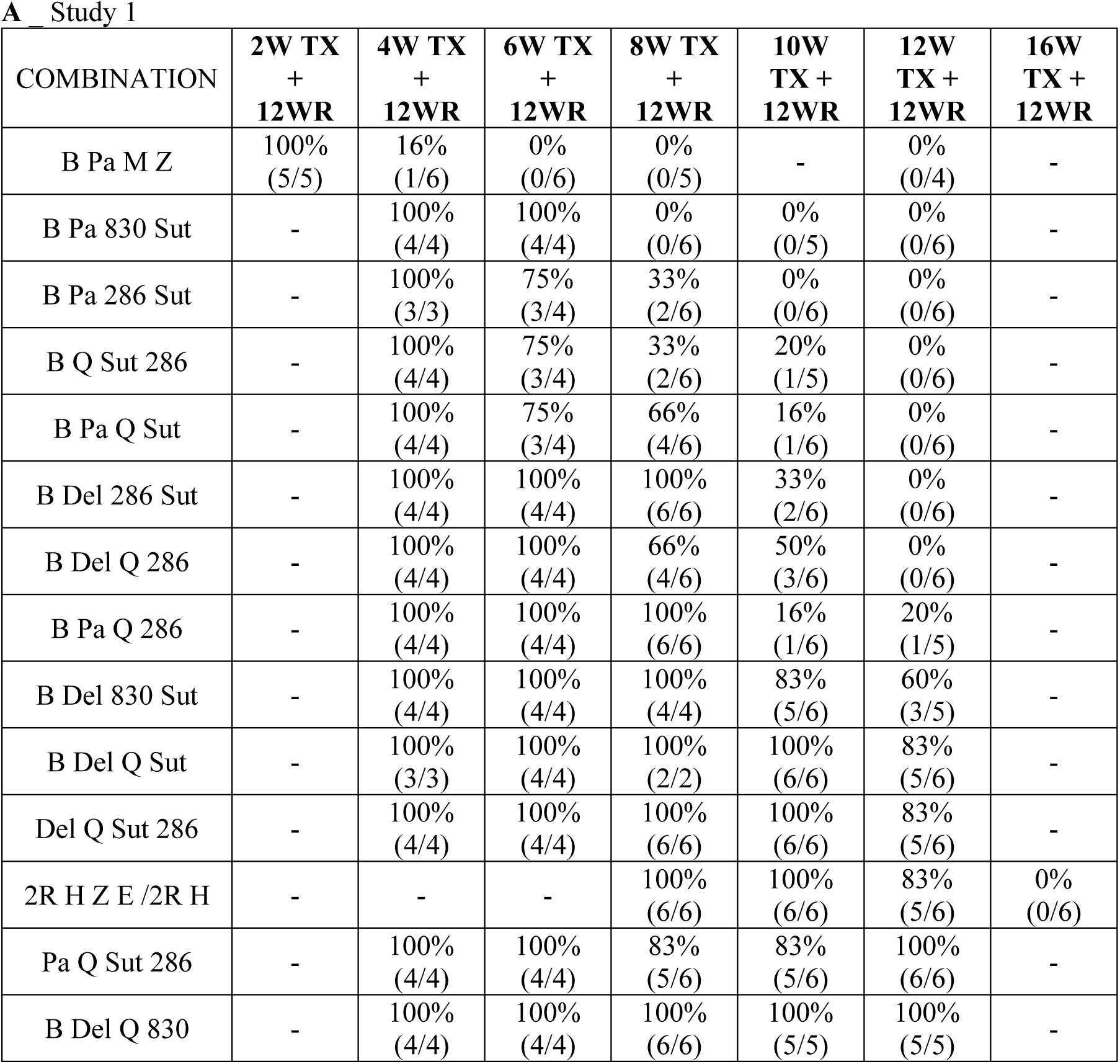
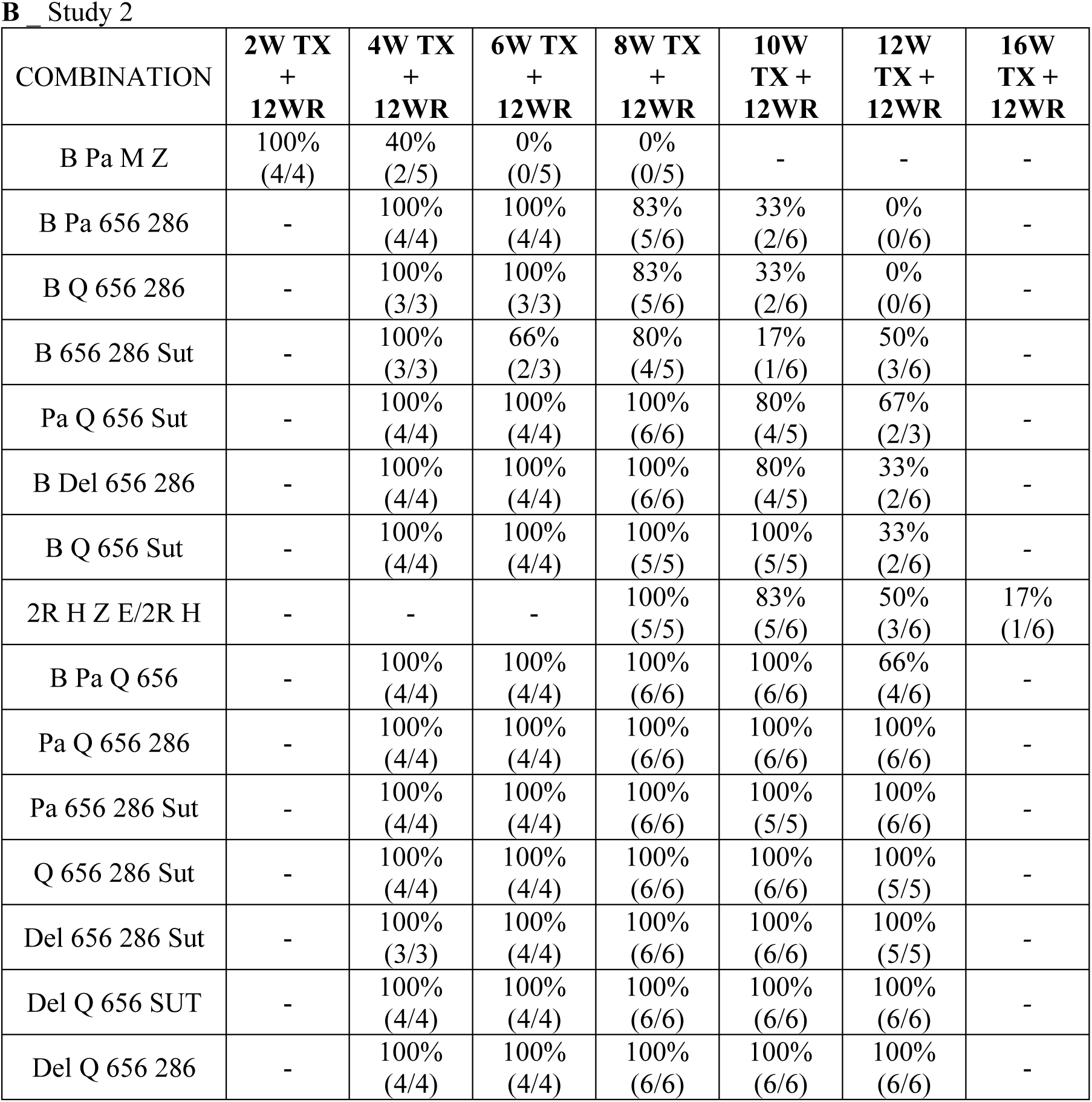
Percentage of relapsing mice, following different treatment durations with the tested drug combinations in study 1 (A) and in Study 2 (B).

A Study 1
| COMBINATION | 2W TX<br>+<br>12WR | 4W TX<br>+<br>12WR | 6W TX<br>+<br>12WR | 8W TX<br>+<br>12WR | 10W<br>TX +<br>12WR | 12W<br>TX +<br>12WR | 16W<br>TX +<br>12WR |
| --- | --- | --- | --- | --- | --- | --- | --- |
| B Pa M Z | 100%<br>(5/5) | 16%<br>(1/6) | 0%<br>(0/6) | 0%<br>(0/5) | - | 0%<br>(0/4) | - |
| B Pa 830 Sut | - | 100%<br>(4/4) | 100%<br>(4/4) | 0%<br>(0/6) | 0%<br>(0/5) | 0%<br>(0/6) | - |
| B Pa 286 Sut | - | 100%<br>(3/3) | 75%<br>(3/4) | 33%<br>(2/6) | 0%<br>(0/6) | 0%<br>(0/6) | - |
| B Q Sut 286 | - | 100%<br>(4/4) | 75%<br>(3/4) | 33%<br>(2/6) | 20%<br>(1/5) | 0%<br>(0/6) | - |
| B Pa Q Sut | - | 100%<br>(4/4) | 75%<br>(3/4) | 66%<br>(4/6) | 16%<br>(1/6) | 0%<br>(0/6) | - |
| B Del 286 Sut | - | 100%<br>(4/4) | 100%<br>(4/4) | 100%<br>(6/6) | 33%<br>(2/6) | 0%<br>(0/6) | - |
| B Del Q 286 | - | 100%<br>(4/4) | 100%<br>(4/4) | 66%<br>(4/6) | 50%<br>(3/6) | 0%<br>(0/6) | - |
| B Pa Q 286 | - | 100%<br>(4/4) | 100%<br>(4/4) | 100%<br>(6/6) | 16%<br>(1/6) | 20%<br>(1/5) | - |
| B Del 830 Sut | - | 100%<br>(4/4) | 100%<br>(4/4) | 100%<br>(4/4) | 83%<br>(5/6) | 60%<br>(3/5) | - |
| B Del Q Sut | - | 100%<br>(3/3) | 100%<br>(4/4) | 100%<br>(2/2) | 100%<br>(6/6) | 83%<br>(5/6) | - |
| Del Q Sut 286 | - | 100%<br>(4/4) | 100%<br>(4/4) | 100%<br>(6/6) | 100%<br>(6/6) | 83%<br>(5/6) | - |
| 2R H Z E /2R H | - | - | - | 100%<br>(6/6) | 100%<br>(6/6) | 83%<br>(5/6) | 0%<br>(0/6) |
| Pa Q Sut 286 | - | 100%<br>(4/4) | 100%<br>(4/4) | 83%<br>(5/6) | 83%<br>(5/6) | 100%<br>(6/6) | - |
| B Del Q 830 | - | 100%<br>(4/4) | 100%<br>(4/4) | 100%<br>(6/6) | 100%<br>(5/5) | 100%<br>(5/5) | - |

**Table 4** Percentage of relapsing mice, following different treatment durations with the 878 tested drug combinations in study 1 (A) and in Study 2 (B).
| COMBINATION | 2W TX<br>+<br>12WR | 4W TX<br>+<br>12WR | 6W TX<br>+<br>12WR | 8W TX<br>+<br>12WR | 10W<br>TX +<br>12WR | 12W<br>TX +<br>12WR | 16W<br>TX +<br>12WR |
| --- | --- | --- | --- | --- | --- | --- | --- |
| B Pa M Z | 100%<br>(4/4) | 40%<br>(2/5) | 0%<br>(0/5) | 0%<br>(0/5) | - | - | - |
| B Pa 656 286 | - | 100%<br>(4/4) | 100%<br>(4/4) | 83%<br>(5/6) | 33%<br>(2/6) | 0%<br>(0/6) | - |
| B Q 656 286 | - | 100%<br>(3/3) | 100%<br>(3/3) | 83%<br>(5/6) | 33%<br>(2/6) | 0%<br>(0/6) | - |
| B 656 286 Sut | - | 100%<br>(3/3) | 66%<br>(2/3) | 80%<br>(4/5) | 17%<br>(1/6) | 50%<br>(3/6) | - |
| Pa Q 656 Sut | - | 100%<br>(4/4) | 100%<br>(4/4) | 100%<br>(6/6) | 80%<br>(4/5) | 67%<br>(2/3) | - |
| B Del 656 286 | - | 100%<br>(4/4) | 100%<br>(4/4) | 100%<br>(6/6) | 80%<br>(4/5) | 33%<br>(2/6) | - |
| B Q 656 Sut | - | 100%<br>(4/4) | 100%<br>(4/4) | 100%<br>(5/5) | 100%<br>(5/5) | 33%<br>(2/6) | - |
| 2R H Z E/2R H | - | - | - | 100%<br>(5/5) | 83%<br>(5/6) | 50%<br>(3/6) | 17%<br>(1/6) |
| B Pa Q 656 | - | 100%<br>(4/4) | 100%<br>(4/4) | 100%<br>(6/6) | 100%<br>(6/6) | 66%<br>(4/6) | - |
| Pa Q 656 286 | - | 100%<br>(4/4) | 100%<br>(4/4) | 100%<br>(6/6) | 100%<br>(6/6) | 100%<br>(6/6) | - |
| Pa 656 286 Sut | - | 100%<br>(4/4) | 100%<br>(4/4) | 100%<br>(6/6) | 100%<br>(5/5) | 100%<br>(6/6) | - |
| Q 656 286 Sut | - | 100%<br>(4/4) | 100%<br>(4/4) | 100%<br>(6/6) | 100%<br>(6/6) | 100%<br>(5/5) | - |
| Del 656 286 Sut | - | 100%<br>(3/3) | 100%<br>(4/4) | 100%<br>(6/6) | 100%<br>(6/6) | 100%<br>(5/5) | - |
| Del Q 656 SUT | - | 100%<br>(4/4) | 100%<br>(4/4) | 100%<br>(6/6) | 100%<br>(6/6) | 100%<br>(6/6) | - |
| Del Q 656 286 | - | 100%<br>(4/4) | 100%<br>(4/4) | 100%<br>(6/6) | 100%<br>(6/6) | 100%<br>(6/6) | - |

A logistic Emax model was developed based on observed cure/relapse data reported herein plus a historical dataset and used to estimate time-to-cure 50% of mice (T50), and time-to- cure 90% of mice (T90) for each combination [23, 37]. Combinations for which 100% relapse was still observed after the maximum treatment duration were excluded. The developed model, using a population approach, demonstrated generally good performance in predicting relapse profiles of the 4-drug combinations, especially for comparators BPaMZ and RHZE/RH for which data coming from different studies and labs were available (**Figure S9**). Individual probability of relapse-time curves for each combination by study are shown in **Figure 3**. The corresponding estimated population T50 and derived T90 values are listed in **Table 5**. T90 values for the 17 analyzed novel combinations ranged from 2 to 6 months, whereas the T90 for the RHZE/RH comparator was approximately 5 months. Fifteen combinations performed better than RHZE/RH, with shorter T90s (**Table 5** and **Figure 3**). Of these, BDel286Sut, BPa656286, BQ656286, BPaQSut, BDelQ286, BPaQ286 displayed a T90 of less than 4 months. The best-performing novel combinations, BPa830Sut, BPa286Sut, and BQSut286, had a derived population T90 less than 3 months with values of 2.11, 2.51 and 2.94 months respectively. None of the test combinations had population time-to-cure parameter estimates T50 or T90 that were as low as for BPaMZ (T90 of approximately one month). When compared to population derived T90s for the combinations included in the historical dataset, some novel PAN-TB combinations appeared to have out-performed clinical reference combinations in this BALB/c mouse model (**Figure 4)**. For example, BPa830Sut and BPa286Sut displayed T90s similar to BPaL (2.28 months) which is used for treatment of adults with Extensively Drug-resistant TB (XDR-TB) or drug-intolerant or non-responsive Multi-Drug-Resistant TB (MDR-TB) using a 6-month regimen [6] (**Table 5 and Figure 4**).

**Figure 3.** Probability of relapse with treatment duration for BALB/c mice treated with combinations tested in Study 1 and Study 2 RMM studies (ordered by individual T90 in the legend). Sterilization curves indicating the probability of relapse over treatment time, constructed by fitting observed relapse data (Table 4) to an Emax model developed using a large historical RMM dataset. Observed relapse data are indicated for each test regimen using open symbols and crosses. The time to 50% cure/relapse is estimated from these curves, and the time to 90% cure (i.e. 10% relapse) is derived utilizing time to 50% cure estimates together with steepness of the curve (gamma) as explained in Methods.

**Figure 4.** Ranking of population T90 values with 95% confidence intervals for all combinations (historical dataset + Study 1 and Study 2 RMM combinations).

**Table 5.** Estimated population T50 and derived population T90 values for combinations tested in Study1 and Study 2 RMM studies. NC, Not computable for combinations which showed 100% relapse at the last treatment time point.

| STUDY | Combinations | Estimated<br>Population<br>T50 (Months) | Derived<br>Population<br>T90 (Months) |
| --- | --- | --- | --- |
| **Both** | **B Pa M Z** | **0.86** | **1.06** |
| Study 1 | B Pa 830 Sut | 1.95 | 2.11 |
| Study 1 | B Pa 286 Sut | 2.04 | 2.51 |
| Study 1 | B Q Sut 286 | 2.11 | 2.94 |
| Study 1 | B Del 286 Sut | 2.97 | 3.22 |
| Study 2 | B Pa 656 286 | 2.85 | 3.38 |
| Study 2 | B Q 656 286 | 2.85 | 3.39 |
| Study 1 | B Pa Q Sut | 2.32 | 3.48 |
| Study 1 | B Del Q 286 | 2.77 | 3.56 |
| Study 1 | B Pa Q 286 | 2.92 | 4.00 |
| Study 2 | B Q 656 Sut | 3.74 | 4.04 |
| Study 2 | B Del 656 286 | 3.60 | 4.28 |
| Study 2 | B Pa Q 656 | 3.99 | 4.40 |
| Study 1 | Del Q Sut 286 | 4.13 | 4.60 |
| Study 1 | B Del Q Sut | 4.13 | 4.60 |
| Study 1 | B Del 830 Sut | 3.93 | 5.08 |
| **Both** | **R H Z E / R H** | **4.15** | **5.09** |
| Study 2 | Pa Q 656 Sut | 4.08 | 5.35 |
| Study 2 | B 656 286 Sut | 2.92 | 6.07 |
| Study 1 | Pa Q Sut 286,<br>B Del Q 830 | NC | NC |
| Study 2 | Q 656 286 Sut,<br>Del Q 656 Sut,<br>Pa Q 656 286,<br>Del 656 286 Sut,<br>Del Q 656 286,<br>Pa 656 286 Sut | NC | NC |

### Comparison of T90s for combinations differing by one drug, indicate treatment-shortening contributions for several PAN-TB candidates, that may be context dependent

Pairwise comparisons of T90 values were performed for 4-drug combinations differing in only one component. In all cases, the presence of B reduced the T90 values and among evaluable comparisons, the T90 differences were statistically significant (p < 0.05) (**Table 6 and Table S3**). For combinations differing only in the nitroimidazole (Pa versus Del), several that contained Pa had a shorter population derived T90 value, compared to those containing Del: Significant differences in T90s were observed for BPa830Sut vs BDel830Sut (2.1 vs 5.1 months); BPaQSut vs BDelQSut (3.5 vs 4.6 months); BPa656286 vs BDel656286 (3.4 vs 4.3 months); and BPa286Sut vs BDel286Sut (2.5 vs 3.2 months). (**Table 6 and Table S3**). However, exposures of B and Q are about 2-times higher in Pa containing regimens than those of in the Del-containing regimens. Therefore, this difference between Pa and Del may be partly explained by the different drug-drug interactions in this mouse model. For certain 4-drug combinations population T90 values did not differ when Q replaced a nitroimidazole: no significant differences or a low level of confidence in the difference were observed between T90s for BQSut286 vs BDel286Sut (2.9 vs 3.2 months) or BQ656286 vs BPa656286 (3.4 vs 3.4 months) or BQSut286 vs BPa286Sut (2.9 vs 2.5 months). On the other hand, replacing Del with Q significantly reduced the T90 for BDel656286 vs BQ656286 (4.3 vs 3.4 months) (**Table 6 and Table S3**).

**Table 6.** Statistical comparison of T90 and AUC_ratio_ between combinations differing by one drug. Green (Delta T90 < 0) and red (Delta T90 > 0) indicate statistically significant differences (p < 0.05) in T90 between Combination 1 and Combination 2. Green (AUC_ratio_ >2; AUC_0-24_ Drug_Combination1_ > 2×AUC_0-24_ Drug_Combination2_) and red (AUC_ratio_ <0.5; AUC_0-24_ Drug_Combination1_ < 0.5×AUC_0-24_ Drug_Combination2_) indicate statistically significant differences in AUC_ratio_. Grey cells indicate not available delta T90 due to not computable T90 values (assumed > 3 months). NS means “Not significant” (p > 0.05). NC means “Not Computable” T90 value. 830 and 656 are considered as equivalent compounds.

| Drugs difference | Comparison (Comb1 vs. Comb2) | Delta T90 | AUC<sub>ratio</sub> Drug 1 | AUC<sub>ratio</sub> Drug 2 | AUC<sub>ratio</sub> Drug 3 |
| --- | --- | --- | --- | --- | --- |
| **656 - 286** | BPaQ656 vs. BPaQ286 | NS | B | Pa | Q |
| **830 - 286** | BDeIQ830 vs. BDeIQ286 | > 2 months | B | Del | Q |
|  | BDeI830Sut vs. BDeI286Sut |  | B | Del | Sut |
|  | BPa830Sut vs. BPa286Sut |  | B | Pa | Sut |
| **B - Del** | BQ656286 vs. DelQ656286 | < -2 months | Q | 656 and/or 830 | 286 |
|  | BQ656Sut vs. DelQ656Sut | < -1 months | Q | 656 and/or 830 | Sut |
|  | BQSut286 vs. DelQSut286 |  | Q | Sut | 286 |
|  | B656286Sut vs. Del656286Sut | < 1 months | 656 and/or 830 | 286 | Sut |
| **B - Pa** | BQ656286 vs. PaQ656286 | < -2 months | Q | 656 and/or 830 | 286 |
|  | BQ656Sut vs. PaQ656Sut |  | Q | 656 and/or 830 | Sut |
|  | BQSut286 vs. PaQSut286 | < -3 months | Q | Sut | 286 |
|  | B656286Sut vs. Pa656286Sut | < 1 months | 656 and/or 830 | 286 | Sut |
| **B - Q** | B656286Sut vs. Q656286Sut | < 1 months | 656 and/or 830 | 286 | Sut |
| **Del - 656** | BDeI286Sut vs. B656286Sut |  | B | 286 | Sut |
| **Del - Q** | BDeI656286 vs. BQ656286 |  | B | 656 and/or 830 | 286 |
|  | BDeI830Sut vs. BQ656Sut |  | B | 656 and/or 830 | Sut |
|  | BDeI286Sut vs. BQSut286 | NS | B | 286 | Sut |
|  | Del656286Sut vs. Q656286Sut | NC | 656 and/or 830 | 286 | Sut |
| **Pa - 656** | BPa286Sut vs. B656286Sut |  | B | 286 | Sut |
| **Pa - Del** | BPaQ656 vs. BDeIQ830 | < -1 months | B | Q | 656 and/or 830 |
|  | BPaQ286 vs. BDeIQ286 | NS | B | Q | 286 |
|  | BPaQSut vs. BDeIQSut |  | B | Q | Sut |
|  | BPa656286 vs. BDeI656286 |  | B | 656 and/or 830 | 286 |
|  | BPa830Sut vs. BDeI830Sut |  | B | 656 and/or 830 | Sut |
|  | BPa286Sut vs. BDeI286Sut |  | B | 286 | Sut |
|  | PaQ656286 vs. DelQ656286 | NC | Q | 656 and/or 830 | 286 |
|  | PaQ656Sut vs. DelQ656Sut | < 0 months | Q | 656 and/or 830 | Sut |
|  | PaQSut286 vs. DelQSut286 | > 1 months | Q | Sut | 286 |
|  | Pa656286Sut vs. Del656286Sut | NC | 656 and/or 830 | 286 | Sut |
| **Pa - Q** | BPa656286 vs. BQ656286 | NS | B | 656 and/or 830 | 286 |
|  | BPa830Sut vs. BQ656Sut |  | B | 656 and/or 830 | Sut |
|  | BPa286Sut vs. BQSut286 |  | B | 286 | Sut |
|  | Pa656286Sut vs. Q656286Sut | NC | 656 and/or 830 | 286 | Sut |
| **Q - 286** | BDeIQSut vs. BDeI286Sut |  | B | Del | Sut |
|  | BPaQSut vs. BPa286Sut |  | B | Pa | Sut |
| **Q - 656** | BDeIQ286 vs. BDeI656286 |  | B | Del | 286 |
|  | BPaQ286 vs. BPa656286 | NS | B | Pa | 286 |
| **Q - 830** | BDeIQSut vs. BDeI830Sut | NS | B | Del | Sut |
|  | BPaQSut vs. BPa830Sut |  | B | Pa | Sut |
| **Sut - 286** | BDeIQSut vs. BDeIQ286 |  | B | Del | Q |
|  | BDeI830Sut vs. BDeI656286 |  | B | Del | 656 and/or 830 |
|  | DelQ656Sut vs. DelQ656286 | NC | Del | Q | 656 and/or 830 |
|  | BPaQSut vs. BPaQ286 | NS | B | Pa | Q |
|  | BPa830Sut vs. BPa656286 |  | B | Pa | 656 and/or 830 |
|  | PaQ656Sut vs. PaQ656286 | < 0 months | Pa | Q | 656 and/or 830 |
|  | BQ656Sut vs. BQ656286 |  | B | Q | 656 and/or 830 |
| **Sut - 656** | DelQSut286 vs. DelQ656286 | < -1 months | Del | Q | 286 |
|  | BPaQSut vs. BPaQ656 |  | B | Pa | Q |
|  | PaQSut286 vs. PaQ656286 | NC | Pa | Q | 286 |
|  | BQSut286 vs. BQ656286 |  | B | Q | 286 |
| **Sut - 830** | BDeIQSut vs. BDeIQ830 | < -1 months | B | Del | Q |

The presence of 286 had a variable impact on 4-drug combination T90 depending on the companion drugs. A reduction in T90 (faster cure) was observed when 286 replaced Q in all tested regimens (BDel286Sut vs BDelQSut (3.2 vs 4.6 months) or BPa286Sut vs BPaQSut (2.5 vs 3.5 months)), and when 286 replaced 830, 656 or Sut in some tested regimens: (BDel286Sut vs BDel830Sut (3.2 vs 5.1 months); BQ656286 vs BQ656Sut (3.4 vs 4.0 months); BDelQ286 vs BDelQSut (3.6 vs 4.6 months) and BDel656286 vs BDel830Sut (4.3 vs 5.1 months)). Finally, the presence of Sut significantly reduced (improved) the T90 when it replaced 656 (BPaQSut vs BPaQ656 (3.5 vs 4.4 months) or BQSut286 vs BQ656286 (2.9 vs 3.4 months). As for B, Sut was present in all 3 best-performing 4-drug combinations that displayed the shortest T90s: BPa830Sut, BPa286Sut and BQSut286.

In addition to these T90 pairwise comparisons, AUC_0-24_ ratios (AUC_ratio_) were calculated and compared for each of the 3 common drugs between pairs of combinations that differed by only one drug. Overall, when only one drug differed, exposure ratios for the common drugs in paired combinations were variable -, although some significant ratios were observed (p < 0.001) very few are more than 2-fold. (**Table 6 and Table S3**).

## DISCUSSION

This translational research utilized the well-established BALB/c TB RMM within a modelling-based framework to evaluate and compare bactericidal and sterilizing efficacy of 25 novel 4-drug combinations of 8 PAN-TB candidate drugs, administered at doses targeting clinically-relevant exposures. The absolute and relative time-dependent bactericidal activities of the RHZE/RH and BPaMZ regimens, as observed here, were consistent with previous data published by Evotec and others [18, 19, 20]. Time-dependent sterilizing activity, evaluated by modelling BALB/c RMM data, has been estimated using varying methodologies by several groups. Despite these differing approaches, the derived population T90s for RHZE/RH and BPaMZ in the present study are consistent with published values [21, 22, 23]. We identified 15 novel PAN-TB 4-drug combinations - i.e. with relevance to the WHO pan-TB Target Regimen Profile - with shorter time to cure (population derived T90) in this model than the current standard of care for DS-TB, RHZE/RH.

The best-performing combinations, BPa830Sut, BPa286Sut, and BQSut286, achieved T90s of less than 3 months in this BALB/c TB model and for BPa830Sut and BPa286Sut, T90 values were similar to BPaL. This means that if developable and safe, these 2 regimens could represent alternatives to BPaL with a pan-TB potential.

None of the PAN-TB combinations cured mice as rapidly as the benchmark regimen BPaMZ, which demonstrated potential to cure patients in less time than RHZE/RH in clinical trials. However, it is promising that several novel 4-drug combinations with pan-TB potential cured BALB/c mice faster than RHZE/RH and at least as rapidly as BPaL. Further, both BPa286Sut and BQSut286 demonstrated rapid bactericidal activity, similar to that of BPaMZ. Rapidly bactericidal regimens may be beneficial by quickly reducing symptoms and infectiousness of TB patients, positively influencing patient and population level outcomes. The observation that BPa286Sut and BQSut286 did not match the cure rate of BPaMZ even after demonstrating similar bactericidal activity is interesting and deserves further attention. First, it underscores what has always been known in TB therapy that rapid elimination of replicating organisms does not necessarily lead to rapid sterilization and lasting cure, even for regimens that contain highly efficacious and long half-life agents like B. Second, it points to a certain sterilizing activity for M and Z, the two agents in BPaMZ, and demands a concerted effort in the field to find agents with similar sterilizing potential.

All 10 PAN-TB combinations that demonstrated T90s of less than 4 months contained B - including the 5 with the most rapid bactericidal activity. In contrast, most B-free combinations didn’t clear lung bacteria after 8 weeks’ treatment or cure mice after 12 weeks of treatment. Both these findings highlight the potentially significant contribution of B and possibly other diarylquinolines to pan-TB treatments. This is in line with previous work demonstrating the importance of this ATP-synthase inhibitor class in the TB treatment backbone [13, 19, 24]. The introduction of B has significantly improved treatment outcomes and survival among patients with MDR/XDR-TB. However, since its implementation in 2012, reports of B resistance have emerged [26]. The identification of potent B-free regimens is therefore of interest. The novel B-free 4-drug combination DelQSut286 demonstrated a slightly shorter T90 than RHZE/RH (by 0.4 months) in this study, offering a qualified hope for B-free regimen for the future.

These results also confirmed the contribution of protein synthesis inhibitors to the performance of novel regimens, with the oxazolidinone Sut present in all 3 combinations with T90s of less than 3 months. Indeed, interest in Sut as a potential novel TB drug led to its evaluation as part of regimens in the Phase2b SUDOCU trial [39] and the PAN-TB Gates MRI-TBD06-21-trial (Int J Tuberc Lung Dis 2025: 29 (11 Suppl 1): S1 – S963). Further, the novel drug candidate 286, which inhibits cholesterol metabolism [3] - a new mode of action for potential TB drugs - is included in 2 of the top 3. The sterilizing activity observed here for BPa286Sut is similar to published data for BPa286L, where full sterilization of mice was achieved within 3 months [12]. L, like Sut, is an oxazolidinone drug and has demonstrated treatment-shortening activity in mice [16, 17]. Notably, in the study of Nuermberger et al [12], undetectable levels of mouse lung CFU were reached after 2 months of treatment with BPa286L as opposed to 4 weeks for BPa286Sut in our study. This difference in bactericidal kinetics may be related to a more potent bactericidal activity of Sut, compared to L, as shown in the study of Tasneen et al [24]. In addition to Sut, the oxaborole leucyl tRNA synthetase inhibitor 830 was present in one of the top 3 regimens, demonstrating the potential utility of this novel mechanism of action which also inhibits protein synthesis, for the treatment of TB.

This is the first time preclinical studies have compared head-to-head bactericidal and sterilizing activity for drug combinations including the two nitroimidazole drugs Pa and Del, at doses confirmed to be relevant to their respective clinical exposures. In these studies, Pa was dosed at 40 mg/kg and Del at 2 mg/kg, lower doses than used in some previously published BALB/c RMM studies [13, 14, 27]. Overall, and especially when considering T90s, the Pa-containing combinations tested here performed better than those containing Del. However, exposures of B and Q are about 2-times higher in Pa containing regimens than those of in the Del-containing regimens. Since B is a known potent sterilizing drug, this difference between Pa- and Del-containing regimens may be partly explained by the different drug-drug interactions in this mouse model. Whether similar drug-drug interactions occur in humans will need to be carefully evaluated. In previous work conducted to compare their bactericidal activity as monotherapies in two mouse models, Del at 2.5 mg/kg demonstrated similar efficacy to Pa at 20 mg/kg and 30 mg/kg [22]. Our results are consistent with this outcome, taking into account the higher dose and presumably exposure of Pa used in our studies. Interestingly, the difference observed in sterilizing activity between Pa-containing and Del-containing matched combinations was sometimes but not always observed when comparing their relative bactericidal activity. For example, whereas BPa286Sut and BDel286Sut reduced lung CFU to undetectable levels at the end of 4 and 8 weeks’ treatment respectively, with corresponding T90s of 2.5 vs 3.2 months, this correlation between bactericidal and sterilizing activities was not observed for BPa830Sut and BDel830Sut. Both regimens reduced bacterial levels to undetectable levels with 8 weeks’ treatment, whereas there is a 3-month difference in T90 between the two combinations (T90s of 2.1 and 5.1 months respectively). Further work is needed to understand the drivers of sterilizing versus bactericidal activity for these combinations and to identify the PK and/or PD factors driving the differentiation in behavior between these two sets of nitroimidazole-containing combinations. Some possible contributors are greater post-antibiotic effect (PAE) for Pa than for Del [28], differentiated PD interactions with companion drugs for Pa versus Del, and differing impacts on exposures of highly efficacious companion drugs for the two nitroimidazoles.

Our pop-PK analysis indicated that differences in T90s were associated with significantly lower AUC_0-24_ values for B or Q, or both, in Del-containing combinations compared to the Pa-containing equivalents while exposure was in the 2x range of clinical target for almost all the comparisons. B exposure in BDelQ830 is the lowest compared to all other B-containing combinations and significantly lower (more than 2-fold) compared to BPaQ656. (**Table 6, Figure S4, Table S3**). In other hand, this analysis indicated that differences in differences in time to cure were associated with a significant lower AUC_0-24_ values for Q in all B-free combinations compared to the B- containing combinations (**Table 6, Table S3**). More generally, numerical results, included in **Table S3**, suggest variable relationships between drug exposure variations and T90 changes. Although some statistically significant AUC_ratio_ were observed, most of them are not clinically relevant as they are within the 2-fold range of clinical target, it is difficult to identify associations between drug exposure and sterilization activity, as significant T90 reductions occur together with either increased or reduced AUC_ratio_ (i.e., >2 or <.0.5, respectively) across paired combinations where one drug differs. Overall, this exploratory analysis highlights that observed exposure-T90 associations must be treated with caution, in the absence of established PK-PD relationships for each compound within the context of these specific 4-drug combinations. Further caution should be exercised when considering translation: any PK drug-drug interaction occurring which may be responsible for observed changes in AUC_ratios_ when one drug is substituted for another, may not translate from mouse to human considering metabolic and/or transporter mechanisms that influence drug (parent and/or metabolite) disposition may differ.

Similar to B-free combinations, nitroimidazole-free combinations are of interest for their utility against nitroimidazole-resistant TB which may emerge in the future [30]. We evaluated the sterilizing activity of the DprE1 inhibitor Q, within multiple 4-drug combinations, for the first time. When either Pa or Del was replaced by Q, these nitroimidazole-free combinations performed as well as the corresponding Pa- containing combinations (i.e. BQSut286; T90 < 3 months) suggesting a similar sterilizing contribution of this DprE1 inhibitor compared to nitroimidazoles, in these particular combinations. The recently reported evaluation of Q together with B and Del in a Phase II clinical trial supports the positive performance of Q within nitroimidazole-containing drug combinations [29]. Our mouse study suggests Q might be useful in nitroimidazole-free regimens

The datasets and modeling approaches used in this work allowed us to compare efficacy of combinations across studies and to historical data, to compare exposures achieved in PD studies to clinical targets and to explore comparative contributions of drugs using PK and PD data. However, further investigation of PK/PD for each agent within the combinations of interest, and direct comparisons of PK and PD for 4-drug combinations versus their 3-drug components, are needed to better understand the contributions of each agent and, in some cases, their active metabolites, to overall combination efficacy, exposure-response relationships and potential PK or PD drug-drug interactions.

These studies constitute the first step in PAN-TB’s nonclinical strategy to prioritize and characterize novel drug combinations to support decision-making and progression through development. The use of this well-established mouse model together with careful dose selection and exposure evaluation, allowed us to assess and compare bactericidal and sterilizing activity of large numbers of drug combinations at relevant exposures in mice as a first step towards prioritization and characterization. The population modelling approaches used for both T90 derivation and PK allowed us to rank combinations with historically tested combinations, across studies and understand the relationship between duration of treatment and cure for each tested combinations while reducing the overall number of mice required. Although the T90 rankings from these BALB/c data appeared generally consistent with clinical results, the model has limitations, including pathology that does not include all lesion types seen in TB patients (e.g those exhibiting caseous necrosis and cavitation).

For this reason, the extent to which findings from BALB/c RMM studies translate to the clinic is hard to predict without further clinical data for comparison to mouse-tested combinations or BALB/c mouse RMM data for regimens recently evaluated in the clinic (e.g Rifapentine (P)HMZ) [31]. Notably, the recently completed Gates MRI-TBD06-201 PAN-TB trial, which evaluated the PAN-TB investigational regimens DBQS and PBQS, concluded that although both regimens demonstrated strong bactericidal activity by the end of treatment when administered for 4 months neither, showed sufficient evidence for being able to treat TB in 3 months or less based on 2- and 3-month sputum culture conversion rates and TB recurrences after treatment completion. When assessed by 12-month post-randomization unfavorable outcome rates, the 4-month PBQS regimen performed similarly to 6-month HRZE (Int J Tuberc Lung Dis 2025: 29 (11 Suppl 1): S1 – S963).

These findings are consistent with the data reported here, where BPaQSut and BDelQSut T90s are 3.48 and 4.6 months, respectively compared with RHZE/RH which cured 90% of mice in 5.09 months.

For combinations of interest identified through these studies, as well as those prioritized based on subsequent BALB/c RMM studies, PAN-TB intends to generate significant additional nonclinical data for use as translational modelling inputs for prediction of clinical performance and to support decision-making. Some examples include nonclinical lesion PK studies, activity studies in caseum (using an ex vivo assay), and evaluation of T90s in mouse models featuring caseous necrotic lesions (the Kramnik RMM). Evaluation of performance of combinations against mouse models where the infecting strain is resistant to a key drug class will also be important. Finally, combination efficacy studies will be conducted in mouse models to better understand the individual contributions of each agent, to confirm their pharmacological value and to inform future regimen design. Promising data has been recently reported for both newer diarylquinolines (e.g. sorfequiline (TBAJ-876) [25]) and oxazolidinones (e.g. TBD09 –[32]) as well as for TBD11, a compound from a different chemical series with a similar mechanism of action as 286, suggest potential to improve on the high-performing first generation PAN-TB regimens by introducing these next generation compounds. Accordingly, the PAN-TB collaboration has introduced sorfequiline, TBD09 and TBD11 to its candidate pool and in addition to seeking a better understanding of the top performing regimens reported here, is conducting further BALB/c RMM studies using the above workflow, to evaluate and compare their bactericidal and sterilizing activities at exposures likely to be targeted in the clinic in potential next generation regimens. The most important outcome of this first study from the PAN-TB consortium has been the successful assembly of novel drugs from different partner organizations and designing and testing drug combinations in a consistent format that will highlight the most promising regimens ready for direct clinical evaluation, thus reducing the time to delivery of these needed interventions to patients.

## MATERIAL AND METHODS

### Animals and ethics

All mouse experiments were carried out at the Evotec France SAS animal facility. This facility is accredited by the French Ministry of Agriculture and by the Association for Assessment and Accreditation of Laboratory Animal Care International (AAALAC). All studies were performed under the European Communities Council Directive (2010/063/EU) for the care and use of laboratory animals and approved by local Ethical Committee CEPAL: CE 029 and authorized by the French Ministry of Education, Advanced Studies, and Research. Six-week-old female BALB/cJRj mice from Janvier Laboratories were group housed in bioconfined cages (Isocage, Tecniplast®) under a 12h light:12h dark with free access to filtered water and a standard rodent diet (AO4C, Safe, France). An ambient temperature of 22 ± 2°C, a relative humidity of 55 ± 10 %, and a negative pressure of -20Pa were maintained throughout the study. All mice were allowed to acclimatize to their new environment for at least 5 days prior to the start of the study.

### Drug Formulations and Dosing Strategies

Drugs were acquired or provided by PAN-TB consortium members, and formulations prepared for dosing as follows at the selected doses (**Table 2**). B (J&J IM) was formulated in 20% 2-hydroxypropy-β-cyclodextrin; Pa (TB Alliance) was formulated in 10% Hydroxy-propyl-beta-cyclodextrin and 2% soy lecithin for dosing at 100mg/kg for the BPaMZ control group, and at 40 mg/kg for the PAN-TB 4-drug combinations. M (LTK Laboratories) and Z (Sigma) were co-formulated in water for dosing at 100mg/kg and 150 mg/kg, respectively. R (Sigma) was prepared in water for dosing at 10 mg/kg; H (Sigma), Z (Sigma) and E (Sigma) were co-formulated in water for dosing at 10, 150 and 100mg/kg, respectively. Del (Otsuka) and Q (Otsuka) were formulated in 5% Arabic gum. 286 (GSK), 656 (GSK) and 830 (GSK) were formulated in 1% methylcellulose solution. Sut (TB Alliance) was formulated in 0.5% methylcellulose and 5% PEG200. For each PAN-TB combination, each drug was dosed individually with an interval of 2 hours between each drug. The order of dosing for each drug in each combination was based on the half-life of each drug where drugs with longest half-lives were administered first and drugs with shorter half-lives were given later in the day. The drug dosing order for each combination is indicated in the name of the combination i.e. the drug listed first was dosed first, and so on. For the BPaMZ comparator, B was administered first, followed by Pa and finally MZ, dosed as a co-formulation. For the RHZE/RH comparator, R was dosed individually for the first 8 weeks, followed by co-formulated HZE. For the following 8 weeks, R and H were dosed individually. The same 2h interval was applied between 2 administrations for the comparators as for the test combinations.

### Relapsing Mouse Model

The 25 PAN-TB combinations were evaluated across two RMM studies. The first 12 combinations were tested in Study 1 (named Wave 2022 in the C-PATH APEX database: https://c-path.org/tools-platforms/tb-apex) and the second 13 combinations in Study 2 (named Wave 2023 in the C-PATH APEX database: https://c-path.org/tools-platforms/tb-apex). BPaMZ and RHZE/RH comparators were included as controls in each Study. Both studies included 4 to 6 mice per group, per time point, allocated following a similar approach to that reported previously [23] and described in **Table S1**. *M.tb* H37Rv stock solution was prepared at exponential growth phase in 7H9 medium / 10% OADC (oleic acid-albumin dextrose-catalase) / 15% glycerol. At Day -14, female BALB/c mice were anaesthetized with 2.5% isoflurane in 97.5% oxygen and were intranasally infected with 50µL *M.tb* H37Rv at an inoculum level of 4.5 Log_10_ CFU/mouse.. Treatment started 14 days post infection, designated Day 0. All drug combinations except RHZE/RH were dosed once daily by oral gavage for 5 days a week (omitting the weekend) for 2, 4, 6, 8, 10, or 12 weeks. RHZE/RH was dosed for 2, 4, 6, 8, 10, 12, 14 or 16 weeks where RHZE was given for the first 8 weeks followed by RH only for the following 8 weeks. For analysis of mouse lung bacterial burden at the end of each designated treatment period, mice were sacrificed 24h post last dosing. For relapse assessment, mice were sacrificed 12 weeks after the end of each designated treatment period. Following sacrifice, lungs were collected, weighed and lung samples homogenized and plated undiluted, or serially diluted on 7H11-OADC + 0.4% activated charcoal. Plates were incubated at 37°C for 6 weeks for CFU quantification. Undetectable level was considered when no colonies were observed in all plates for the sample.

### Pharmacokinetics

Combination PK studies were conducted, before the RMM studies, in female BALB/c mice, intranasally infected with *M.tb* H37Rv in the same manner as for the RMM studies (see RMM section of Materials and Methods). Beginning 14 days post-infection (Day 0) 24 mice per group were treated with each test combination via oral gavage either once or once daily for 5 days. The 5-day dosing period was selected based on the short terminal half-lives of the drugs (**Table S3**), suggesting at least 80% of the steady-state is reached after 5 days treatment duration (5 days > 5 times of T_1/2_). The only exception is the bedaquiline M2 (desmethyl) metabolite (B-M2), which exhibits a longer half-life. It is assumed that the late sparse PK samples in the RMM study (Week 8) provide enough information to correctly estimate the PK parameters for this entity. Doses were selected based on historical data from PAN-TB members. The order and spacing of administrations were performed as for the RMM studies (see Dose Formulations and Dosing Strategies). Blood samples were collected from the tail vein on days 1 (after single dosing) and 5 (after once daily repeat administration) at 0.5, 2.5, 5.5, 7.5, 10, 12, 14, 24, 36, 29, and 31 hours post administration of the first drug. Additionally, for each combination, plasma and blood intra-cardiac concentrations were assessed on Day 1 and Day 5 at 2.5, 7.5, 14, and 31 hours post-first compound dosing. During RMM studies, on the 5^th^ day of the 8^th^ week of treatment, blood was collected from the tail vein at 0.5, 2.5, 5.5, 7.5, 9 and 24 hours post-first compound dosing. In all cases, after blood processing, each compound, including B-M2, and the active (sulfoxide) metabolite of sutezolid (PNU-101603) were quantified by liquid chromatography tandem mass spectrometry (LC-MS/MS). Bioanalytical methods, samples handling, as well as MS conditions are described in supplemental data. All PK assessments were conducted in blood. Blood and plasma concentrations determined through the Combination PK studies were used to calculate mouse geometric mean Blood-to-Plasma (BP) ratios (**Table 2**) for each drug. These values were used to facilitate translation of blood to plasma exposure values for head-to-head comparisons between observed mouse and clinical target exposures (see Population PK Modelling).

### Population PK Modelling

For each compound tested across the 25 PAN-TB combinations, a population pharmacokinetic (popPK) model was developed to describe the mouse PK profile. After identifying each specific model, post-hoc estimates were used to calculate total exposures, i.e. AUC_0-24_, accounting for the variability observed across animals in terms of summary statistics (median, 5^th^, 25^th^, 75^th^ and 95^th^ percentiles). Blood concentration data collected during the RMMs were pooled with those collected in the Combination PK studies to ensure a sufficient number of observations required for robust model development and parameters estimation. More details about model selection criteria are provided in the supplemental material (see *PopPK model verification and quality criteria*). For B and Sut their main active metabolites, B-M2 and PNU-101603 respectively, were included in the model description to derive the overall exposure. As the parent drug and corresponding metabolite contribute to efficacy in mouse and human at different relative levels [33, 34, 35, 36], the active moiety approach was adopted to compare the relevance of preclinical exposure values to clinical exposures, considering the plasma AUC_0-24_/MIC_90_ ratios (AUC_24MIC_) in both human and mouse:

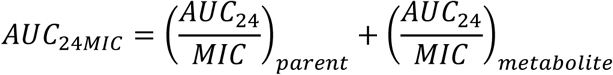

To enable comparisons of mouse exposures against clinical plasma targets, simulated blood concentrations were converted to plasma values using the compound-specific geometric mean Blood-to-Plasma (BP) ratio (**Table 2**). The area under the curve, during the 24 hours post drug administration (AUC_0-24_, ng·h/mL) was determined, assuming a dense simulation time grid (i.e., 0.1 h sampling frequency) which enabled robust determination of the total exposure in plasma. PopPK modelling was performed with Monolix Suite^®^ (Build 2024R1, Lixoft) running on a Windows 11, 64-bit operating system. All models were identified using a Stochastic Approximation Expectation-Maximization (SAEM) algorithm. All analyses for the extrapolation of exposure metrics were performed using R Statistical Software (v4.4.2; R Core Team 2024).

### Logistic Emax Model

A logistic Emax model was applied, as previously described [23] using data obtained from Study 1 and Study 2 plus a historical dataset. The historical dataset consisted of data utilized in previous modelling efforts [37] as well as other literature data [19, 38, 39] and unpublished historical data generated at Evotec. The Evotec internal studies assessed efficacy of RHZ/RH, BPaL and PaMZ dosed 5/7 or 7/7 days per week in a BALB/c RMM and data from these studies is available via the C-PATH APEX platform (https://c-path.org/tools-platforms/tb-apex).

Data were pooled from the same treatment group collected from different studies (i.e. within the historical and present datasets). For each combination, i and study k, γ and T50 parameters were estimated and subsequently used to calculate the related T90, according to the following formula:

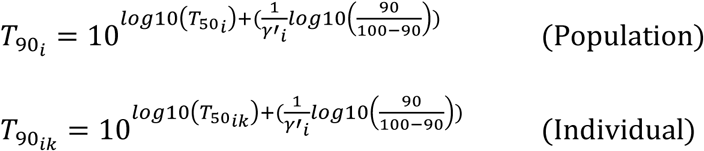

Population parameters T50 and T90 enabled characterization of the average time to 50% or 90% cure respectively, for a particular combination across studies, independent of study-related factors (e.g., starting inoculum), while individual parameters (e.g. T50_ik_ and T90_ik_) provide the average time to 50% or 90% cure for each specific study. To limit the number of parameters to be estimated and overcome identifiability issues, combinations in the historical dataset that share the same drugs but were administered at different dose levels or with differing dose schedules were parametrized with different T50 parameters, assuming the same γ value (i.e. steepness of the relapse-time curve). To estimate model precision and support comparisons of the combinations, confidence intervals (CI) were computed for T90s. Since T90 was derived from T50 and γ estimates, the delta method [41] was employed to compute an approximation of its standard error (SE), assuming that correlation between T50 and γ is negligible (assessed by the non-parametric Spearman correlation test being T50 and γ not normally distributed, with p-value lower than 0.05 considered statistically significant). Once T90 SE had been obtained, 2.5^th^ and 97.5^th^ percentiles (i.e. CI_lower limit_ and CI_upper limit_, respectively) were computed for each combination by assuming normally distributed data, as follows:

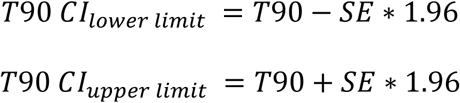

The logistic Emax model was developed using NONMEM® 7.5.1 and data handling was performed using SAS® version 9.4 and R Statistical Software (v4.4.2; R Core Team 2024).

### Statistical Analysis

Statistical analysis was conducted to evaluate differences between a set of pairwise drug combinations that differed by only one component. The analysis considered both overall T90 results and exposure levels (AUC_0-24_) of the drugs common to the combinations. Specifically, T90 results were compared using a Z-test, assuming that T90 values are normally distributed, and their standard errors are known. AUC comparisons were performed using a t-test with unequal variances on log-transformed data. No formal correction for multiple comparisons was applied, as the analyses were exploratory. P-values below 0.05 were considered statistically significant.

## Acknowledgements

This work was supported by the Gates Foundation [INV-008993]. The conclusions and opinions expressed in this work are those of the author(s) alone and shall not be attributed to the Foundation. Under the grant conditions of the Foundation, a Creative Commons Attribution 4.0 License has already been assigned to the Author Accepted Manuscript version that might arise from this submission. Please note works submitted as a preprint have not undergone a peer review process.

We would like to thank the Evotec BSL3 In Vivo Pharmacology, Bioanalytical and Pharmacometrics teams for their valuable support and collaboration. We are especially grateful to Augusto Celon and Andrea Boscolo Panzin for their insightful input and contributions to the analytical aspects of this work, as well as Jeanne Jaen, Fanny Deglave and Eric Erdocain for expert bioanalytical work.

---

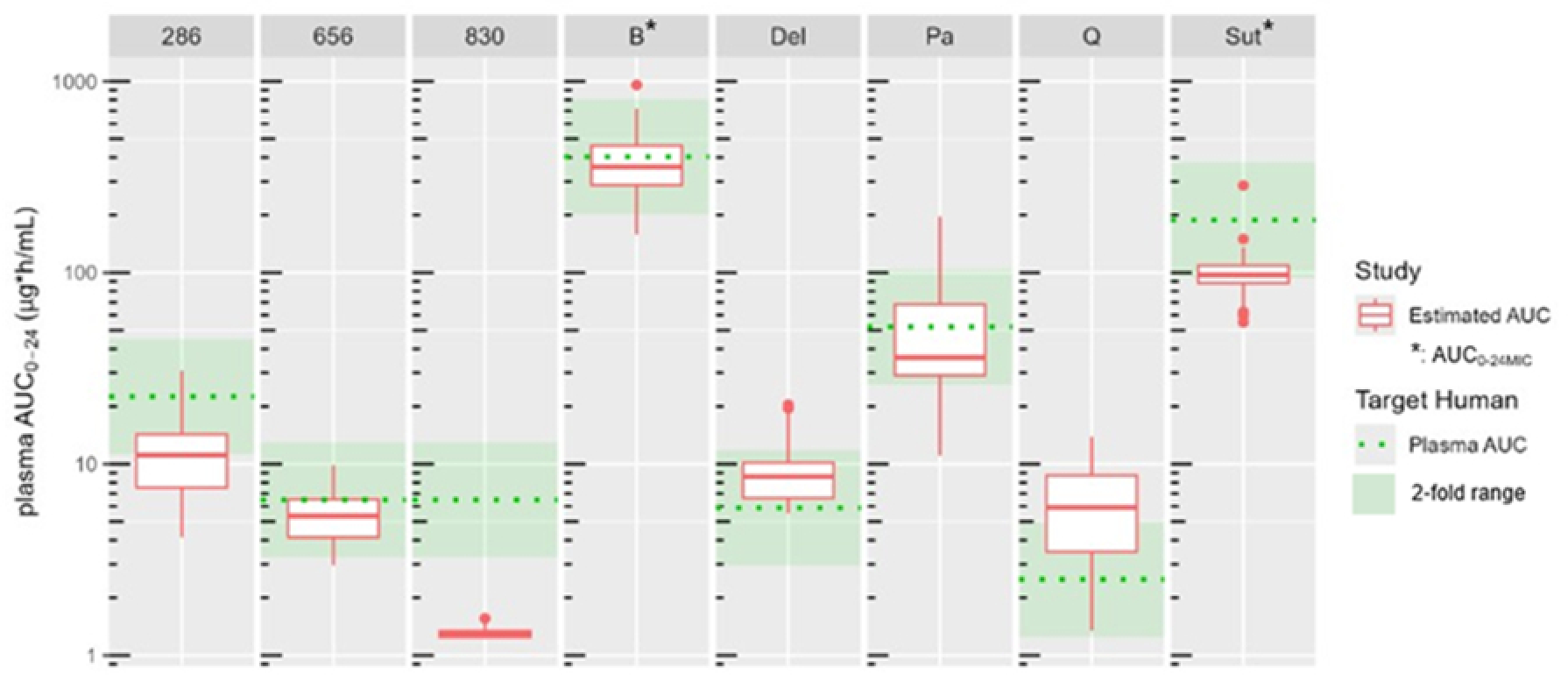

